# A long-context language model for deciphering and generating bacteriophage genomes

**DOI:** 10.1101/2023.12.18.572218

**Authors:** Bin Shao

**Author notes:** Correspondence should be addressed to Bin Shao.

## Abstract

Inspired by the success of large language models, we develop a long-context generative model for genomes. Our multiscale transformer model was pre-trained on unannotated bacteriophage genomes with byte-level tokenization. We demonstrate the foundational capabilities of our model including the prediction of essential genes, genetic variant effects, regulatory element activity and taxonomy of unannotated sequences. Furthermore, it generates *de novo* sequences up to 96K base pairs, which contain functional regulatory elements and novel proteins with phage-related functions.

---

Large pre-trained language models have drastically transformed the natural language processing (NLP) field^1,2^. Drawing on the similarity of natural language and genome sequences, genomic language models have been developed. These models were trained on large scale genomic datasets, and they effectively predict regulatory elements, uncover co-regulation patterns in proteins, and identify the genome-wide variant effects^3–7^. However, it remains an open question whether language models can be tailored to generate genome-scale sequences with functional elements, while retaining the capacity to decipher the intricate relationships within DNA sequences.

Most of the current genomic language models used masked language modeling like Bidirectional Encoder Representations from Transformers (BERT)^1^. This approach is not ideal for tasks that require generating new contents. In addition, they face technical constraints such as short context size and the aggregation of sequences in k-mer tokenization. These limitations hinder their ability to learn from genome-scale data with the level of resolution needed for designing functional elements.

In this work, we introduce megaDNA, a long-context language model that demonstrates foundational capacities in understanding and generating genomic sequences. Our model draws inspiration from the Generative Pre-trained Transformers (GPT) model^2^, which is renowned for its proficiency in generating long and coherent texts. We utilized a multiscale transformer structure^8^ that enables us to train the model on unannotated whole bacteriophage genomes at the single nucleotide-level. Without further training, our model can predict gene essentiality across the phage genome in a zero-shot manner. The model embeddings can be directly applicable to predict functional properties of both regulatory elements and proteins. Moreover, the trained model generates sequences up to 96K base pairs (bp), sharing a similar genomic structure with the natural bacteriophage genomes. We found functional promoters and ribosome binding sites (RBS) in the 5’ untranslated regions (5’UTR) of the predicted genes. The proteins from the generated sequences are predicted to be structurally plausible. Our model is available from GitHub: https://github.com/lingxusb/megaDNA

To construct the training dataset, we collected bacteriophage genomes with high confidence from three sources including the NCBI genebank, the Metagenomic Gut Virus (MGV) catalogue^9^, and the Gut Phage Database (GPD)^10^ (Supplementary Fig. 1). After data cleaning, we constructed a dataset of 99.7K bacteriophage genomes to pre-train our model (Methods). The training data was byte-level tokenized, and we employed a multi-scale transformer structure with three layers and a long-context window in model training, as proposed by Yu et al.^8^

We hypothesize that our pretrained language model captures the structural patterns of bacteriophage genomes, allowing the model’s loss to approximate the fitness of genome sequences. To test this hypothesis, we conducted *in silico* mutagenesis analysis to predict essential genes in the lambda phage genome^11^ (Fig. 1b). Without any supervised training, we found mutations within the coding sequences of essential genes result in higher losses than non-essential genes (Fig. 1c). Consequently, changes in model loss can be used a zero-shot predictor of essential genes (AUROC: 0.86, Fig. 1d). Similarly, mutations in the start and stop codons of essential genes lead to higher model losses than non-essential genes (Fig. 1d, Supplementary Fig. 2).

**Figure 1.**
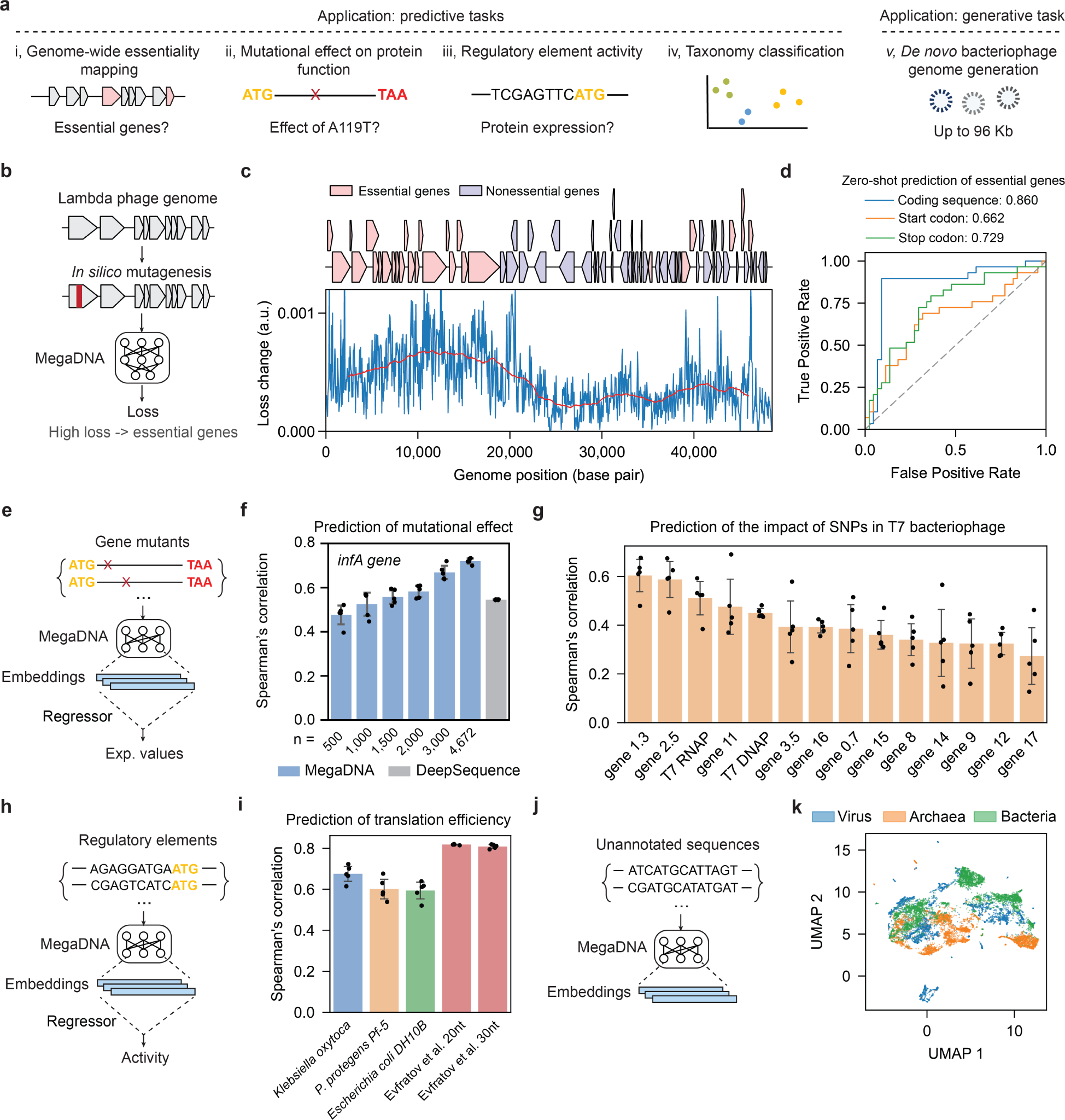
Foundational capacities of our language model. **a**) Overview of the model applications. **b**) *In silico* mutagenesis analysis to identify essential genes in the bacteriophage genome. **c**) Model loss variation across the lambda phage genome in the mutagenesis analysis. Upper, essential and non-essential genes in the genome. Lower: changes in model loss for 50 bp non-overlapping windows across the genome (blue). Moving averages of model loss across 5,000 bp windows are denoted in red. **d**) Zero-shot prediction of essential genes by calculating the effects of mutations in the gene coding region (blue), start codon (orange), and stop codon (green). Area Under the ROC Curve (AUROC) scores are reported. **e**) Prediction of mutational effects on protein functions using model embeddings. **f**) Prediction of mutational effects for the deep mutational scanning experiment of the *infA* gene. Spearman correlation coefficients of the predicted and reported fitness from 5-fold cross validation tests are reported (Blue: megaDNA, gray: DeepSequence). n is the number of training samples. **g**) Prediction of the impacts of Single Nucleotide Polymorphisms (SNPs) in the T7 bacteriophage genome. Spearman correlation of the predicted and reported fitness from 5-fold cross validation tests are reported. **h**) Prediction of regulatory element activity using model embeddings. **i**) Prediction of translation efficiencies for non-model organisms and high-throughput gene expression libraries. For *K. oxytoca, P. protegens* and *E. coli DH10B*, we evaluated the model performance on endogenous genes. Five-Fold cross-validation tests were used for all calculations. **j**) Distinguishing taxonomies of unannotated sequences using model embeddings. **k**) UMAP visualization of model embeddings for sequences from bacteriophages, bacteria, and archaea (model middle layer, n=5000 per group). For f, g and i, error bars denote standard deviation (n = 5).

Sequence embeddings from language models capture rich contextual information. We further explored the utility of our model’s embeddings for a range of predictive tasks. We first evaluated our model’s ability to predict effects of sequence mutations on protein functions (Fig. 1e). Mutated gene coding sequences were used as inputs, and a regressor was trained on model embeddings to predict the mutational effects. Our model’s prediction performance closely matched the state-of-the-art model DeepSequence^12^ (Fig. 1f), including for a protein not existing in the training dataset (Supplementary Fig. 3). Moreover, our model successfully predicted the impact of SNPs across the T7 bacteriophage genome^13^, even with limited training samples (Fig. 1g, Supplementary Fig. 4).

Expression of phage genes relies on the protein synthesis machinery in host bacteria cells. We investigated the potential of the model embeddings to predict regulatory element activity in bacteria (Fig. 1 h). Our model effectively predicted the translation efficiencies of 5’UTR in non-model organisms such as *K. Oxytoca, P. Protegens*, as well as for high-throughput gene expression libraries in *E. coli*^14^ (Fig. 1i). The model performance is also robust to the training sample size (Supplementary Fig. 5).

Lastly, we extended our model to identify taxonomy of unannotated sequences (Fig. 1j). We collected unannotated sequences from bacteriophage, bacteria, and archaea. The embeddings from our model separated these sequences in a low-dimensional space (Fig. 1k). By training a linear regressor based on model embeddings, we achieved a high classification accuracy (average AUROC of 0.98, Supplementary Fig. 6). This high level of accuracy was consistent across different layers of our model (Supplementary Fig. 7). Since the training data doesn’t contain genome sequences of bacteria or archaea, these results demonstrate the broad applicability of our model.

Our approach enables *de novo* generation of genome sequence of bacteriophage (Fig. 2a). We generated a total of 1,024 sequences longer than 1K bp. geNomad^15^ was used for functional annotation of the generated sequences. Among all these sequences, 607 have a virus score larger than zero. The average sequence length is 43K bp, and the average number of predicted genes per sequence is 67, which are similar to the training dataset (mean length: 48K bp, average number of predicted genes: 68). The gene length distribution is close to that of the training dataset (Fig. 1b, average gene length: 558 bp vs 646 bp), while the predicted gene numbers show wider spread (Supplementary Fig. 8). The median virus score of these generated sequences is 0.84 and the maximum score is 0.97, comparable to the virus scores of natural bacteriophage genomes which range from 0.70 to 0.98 (Fig. 1c). 223 out of 607 generated sequences (37%) are predicted to be Caudoviricetes by geNomad (Fig. 1d). As a comparison, 98% of the genomes in the training dataset were classified as Caudoviricetes. Additionally, 388 sequences were predicted to have bacterial hosts with a probability greater than 0.95, as determined by the DeepHost model^16^ (Supplementary Fig. 9).

**Figure 2.**
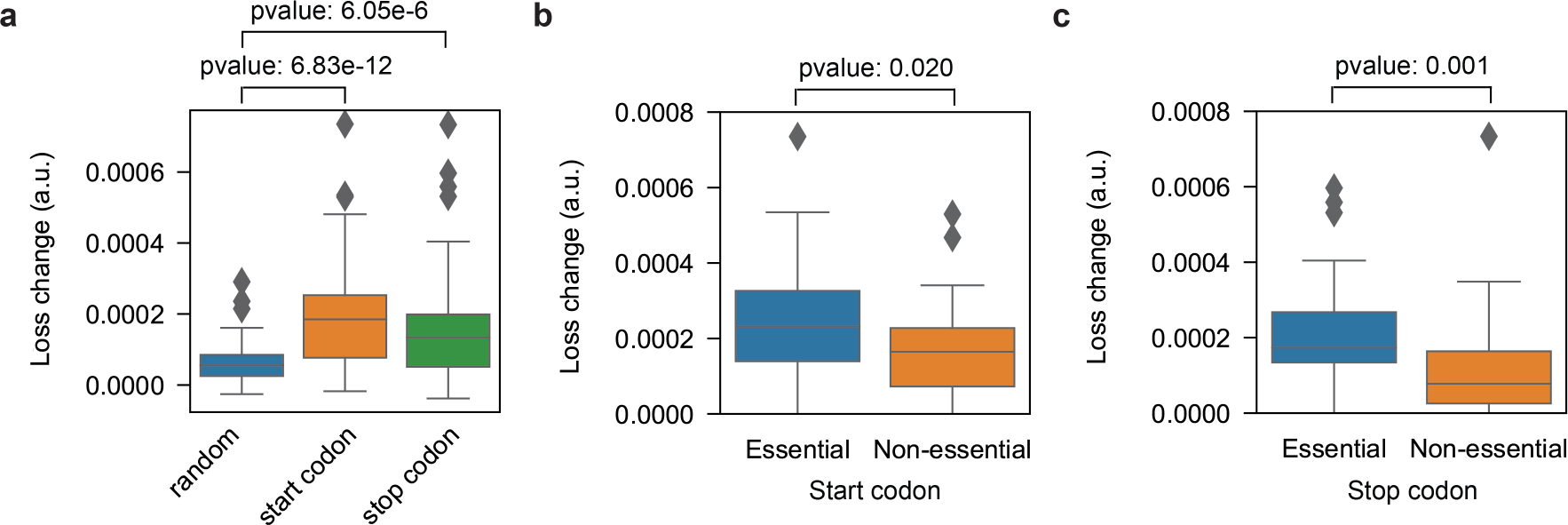
Language model generates sequences with functional genomic structures. **a**) The workflow schematic. **b**) Comparison of gene length distributions between predicted genes in generated sequences (n = 40,399) and a randomly sampled subset of genes from the training dataset (n = 10,000). **c**) Comparison of the predicted virus scores for generated sequences and the training dataset. **d**) Predicted taxonomy for the generated sequences. Only taxonomies with > 1 sequence are shown. **e**) Functional annotation of a selected sequence fragment (generated sequence #87). **f**) Predicted promoter activity for all the 5’UTRs in the generated sequence #87 (n = 44), along with the promoter activity of the random sequences with the same length. **g**) Proportions of adenine (A) and guanine (G) nucleotides preceding the start codon of all the predicted genes in the generated sequence #87. **h**) Mean predicted pLDDT scores for proteins with geNomad markers from generated sequences (sample size: n = 343; median value: 28) against random peptide sequences of the same lengths (sample size: n = 343; median value: 18). A sample generated protein is shown on the right. **j**) Top 10 predicted functions of proteins derived from the generated sequences.

We then examined the 5’UTR of the annotated genes in the generated sequences to determine if they contain functional regulatory elements such as promoters and RBS to initiate transcription and translation. We chose the generated sequence #87 for further analysis due to its high predicted virus score (0.96) and its relatively small size (28K bp). Using a machine learning tool (Promoter Calculator)^17^, we identified the -35 box and -10 box of the promoter within the 5’UTR of the predicted phage stabilization protein. Notably, their sequences are close to the established consensus motifs: TTGACA and TATAAT (Fig. 1f). Prior to the start codon of the same gene, we observed a region enriched in adenine (A) and guanine (G) nucleotides, indicative of a functional ribosome binding site (Fig. 1f).

Analyzing all 5’UTR of the predicted genes from this sequence, we found a significantly higher promoter activity compared to random sequences of the same length (Fig. 1g). Intriguingly, the proportion of A and G nucleotides peaked around 10 bp upstream of the start codon, aligning closely to the optimal position for an RBS to drive translation initiation (Fig. 1h). This trend of A/G enrichment is also consistent across all the generated sequences (Supplementary Fig. 10). In short, our generated sequences harbor functional regulatory sequences that could enable expression of the predicted genes.

In our generated sequences, 343 annotated genes were predicted to match geNomad markers. These genes share very little homology with the training dataset (Supplementary Table 1). We employed ESMfold^18^ to predict their structures and calculated the average predicted local distance difference test (pLDDT) scores. This score reflects the confidence of ESMfold on the predicted structures. The median pLDDT score for these proteins is higher than that of random peptide sequences (28 vs 18). We further randomly sampled 10K annotated genes from all the generated sequences and found high pLDDT scores for them (median value of 36, Supplementary Fig. 11), suggesting that these generated proteins are more likely to adopt a stable conformation. Functional annotation of all annotated proteins using deepFRI^19^ reveals several large families with diverse functional roles, including the transporter activity and the structural molecule activity (Fig. 1j). Among these, several proteins were predicted to have DNA-binding activity, and the predicted structure resembles the canonical helix-turn-helix (HTH) domain in this protein family (Supplementary Fig. 12).

To the best of our knowledge, our work presents the first long-context generative model for genomic sequences. Our language model effectively learns the language of gene coding and regulatory sequences via a single step of self-supervised training on unannotated whole genomes. The generated sequences match the length of natural bacteriophage genomes and display functional genomic architectures. It is worth noting that these sequences have not been optimized at the codon or gene level to allow for efficient self-replication in bacteria. However, with further scaling up and fine-tuning, we envision that generative genomic models have the potential to enable *de novo* design of the whole genome, offering opportunities for breakthroughs in medicine, agriculture, and environmental science. This field also faces ongoing challenges in ethical considerations, biosafety, and regulatory frameworks, which are critical for the responsible advancement of generative modeling in synthetic biology.

## Methods

### Model training

Our training dataset was curated from three sources. Firstly, we downloaded all the complete virus genomes from NCBI genebank, retaining only those with “phage” in the organism’s name. Secondly, the phage genomes from MGV were downloaded, and we only included genomes with a completeness score larger than 95% and classified under the order Caudovirales. Our third source was GPD, and we kept all the genomes with a completeness score above 0.95. Following the initial collection, we undertook an additional round of filtering. We used geNomad to predict the taxonomy of these genomes and then deleted all the genomes whose predicted host is not a unicellular organism. All genomes smaller than 96K bp were used to construct the final training dataset.

Our megaDNA model utilized a three-layer transformer structure^8^. Each layer had a depth of 8 and progressively larger dimensions (local: 196, middle: 256, global: 512). The sequence lengths for three layers are 16, 64, 128. The model contains 145M parameters in total. We assigned numerical tokens (1, 2, 3, and 4) to the nucleotides A, T, C, and G, respectively. For model training, we used a batch size of 1 and set the learning rate at 0.0002. The learning rate was progressively increased during the initial 50,000 steps as part of a warmup schedule. We utilized the Adam optimizer and applied gradient clipping with a norm of 0.5 to prevent gradient explosion.

### *In silico* mutagenesis of phage genomes

Lambda phage genome sequences and annotations were downloaded from NCBI (Accession number: NC_001416.1). Essential genes were identified according to Piya et al^11^. We conducted *in silico* mutagenesis using a 50 bp sliding window across the genome, and each nucleotide was randomly mutated to A, T, C, or G with equal probability. The impact of mutations was assessed by computing the model loss, which is further compared with their original counterparts. For both essential gens and non- essential genes, we calculated the mean model loss for all the windows within the gene coding region. Mann-Whitney U test was used to evaluate statistical differences between these two groups (*scipy.stats.mannwhitneyu*). Furthermore, mutations targeting the start and stop codons of all coding genes were simulated. We also generated a control set comprising an equivalent number of 3-nt mutations randomly distributed across the genome. The effect of these mutations on model loss was analyzed to infer their impact on fitness. The changes in model loss were used a predictor of gene essentiality, and we computed the Receiver Operating Characteristic (ROC) curve and the Area Under the ROC Curve (AUROC).

### Prediction of mutational effects on protein function

The DNA sequences of the mutated genes were used as the model input. Embeddings from three different layers of the model were extracted (dim = 196, 256, 512). These embeddings were concatenated to form a 964-dimensional vector representing each gene coding sequence. We used the 5-fold cross validation to evaluate the prediction performance of our model. In each fold, one fifth of the data was held out as test data while the remaining data were used as training data. A Ridge regression model was trained on the input and fitness values in the training dataset with default parameters (*sklearn.linear_regression.RidgeCV*). The predictive performance of this model was then evaluated on the test dataset. The infA gene dataset was obtained from Kelsic et al^20^. For the T7 bacteriophage dataset^13^, genome sequence and annotations were downloaded from NCBI (Accession number: V01146.1). The model performance was evaluated for each gene in the same manner.

### Prediction of translation efficiency

We assessed the translation efficiency (TE) of genes in *Klebsiella oxytoca, Pseudomonas protegenes Pf-5*, and *Escherichia coli DH10B* by calculating the ratio of average ribosome density (RD) to mRNA expression. The ribosome density of each gene was calculated by averaging all ribosome occupancies over the length of the gene. The mRNA expression in FPKM (fragments per kilobase of transcript per million mapped reads) of each gene that was calculated by averaging the height of RNA-seq profile over the length of the gene. The ribosome profiling and RNA-seq datasets of *K. oxytoca* and *P. protegenes* Pf- 5 were obtained from the Sequence Read Archive with the accession code PRJNA579767^21^. The *E. coli* DH10B datasets were obtained from NCBI GEO database with accession number GSE152664^22^. We used the DNA sequences spanning from -160 to 160 relative to the start codon as the input to our model.

Model embeddings were extracted from three layers (dim = 196, 256, 512) and concatenated together to form a 964-dim vector for each input sequence. To mitigate the influence of lowly expressed genes on TE calculations, we focused on the top 25% expressed genes in *Klebsiella oxytoca* and *Escherichia coli DH10B,* and the top 20% expressed genes in *P. protegenes Pf-5*. We used 5-fold cross validation to evaluate the performance of our model. In each fold, a ridge regression model was trained on the input and TE values in the training dataset with default parameters (*sklearn.linear_regression.RidgeCV*). The trained model was then used to predict TE values in the test dataset. For the Evfratov et al. dataset^14^, 20 nt and 30 nt 5’UTR sequences were used as the input. The model performance was evaluated as previously described.

### Classification taxonomy of unannotated sequences

We analyzed 10K bp sequences randomly sampled from bacteriophage, bacteria, and archaea genomes downloaded from NCBI genebank (n = 5,000 each). For the total of 15,000 sequences, sequence embeddings were generated using our model across three layers (dim = 196, 256, 512). For embedding visualization, we used Uniform Manifold Approximation and Projection (UMAP)^23^ as implemented in the python package *umap-learn*. To classify these sequences, we employed a logistic regression model to evaluate the predictive performance of the embeddings across multiple classes (*sklearn.linear_model.LogisticRegression*). The model’s performance was assessed using a 5-fold Stratified K-Fold cross-validation. This method ensures that each fold is a good representative of the whole by maintaining the same proportion of samples for each class as in the complete dataset. Within each fold, the model was trained on a subset of the data and then used to predict probabilities on the test subset. For each class, we computed the Receiver Operating Characteristic (ROC) curve and the Area Under the ROC Curve (AUROC).

### Model inference

We generated sequences from the trained model using a predefined set of parameters. Specifically, we adjusted the temperature to 0.95 to ensure a balance between variety and coherence in the sequences and kept the filter threshold at 0.0 to avoid limiting the range of token probabilities. For model training and inference, we utilized Nvidia’s A100 GPU (40GB) and 3090 Ti GPU (24GB) and used the PyTorch version 2.1.1 software package.

### Analysis of the generated sequence

geNomad^15^ was used for sequence annotation of all generated sequences. The 100 base pair regions preceding the start codon of each predicted gene was designated as the 5’UTR. We employed the Promoter Calculator^17^ to find the promoters in these regions. Only the promoter with the highest predicted activity was annotated. For protein structure prediction, we used the pretrained ESMfold model v1^18^. The chunk size of the model was set to be 64 for proteins longer than 700 AA and 128 for shorter proteins. We limited our structure calculation to proteins less than 1000 AA in length. Function prediction for these proteins was carried out using the default deepFRI model^19^, as available on GitHub (https://github.com/flatironinstitute/DeepFRI). We used a score cutoff of 0.5 which was reported to be significant in the original publication. Bacteria host prediction was done using DeepHost species-level model^16^ with default parameters (https://github.com/deepomicslab/DeepHost). Predictions with a probability greater than 0.95 were used in the following analysis.

## Data availability

The bacteriophage genomes were downloaded from public databases including NCBI genebank (ftp://ftp.ncbi.nlm.nih.gov/genomes/genbank/), MGV (https://portal.nersc.gov/MGV), and GPD (https://www.sanger.ac.uk/data/gut-phage-database/).

## Code availability

Our trained model and model inference codes are available from GitHub: https://github.com/lingxusb/megaDNA

---

**Supplementary Figure 1:**
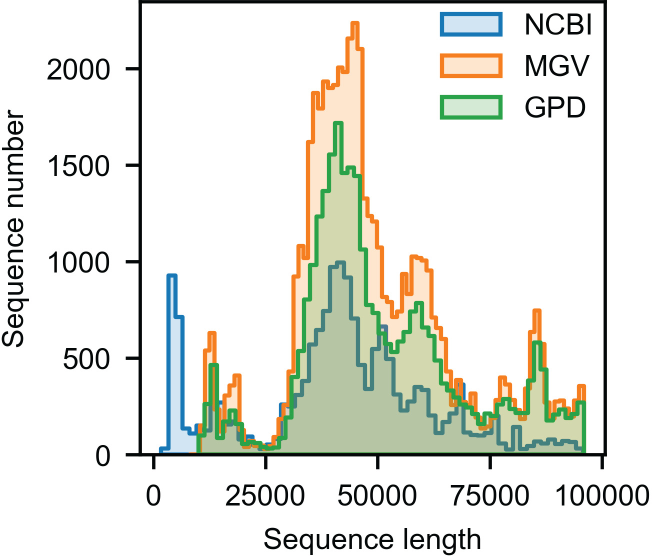
Genome size distributions of three data sources. Distributions of genome sizes within the training dataset: NCBI (n = 16,609), MGV (n = 53,032) and GPD (n = 30,032).

**Supplementary Figure 2:**
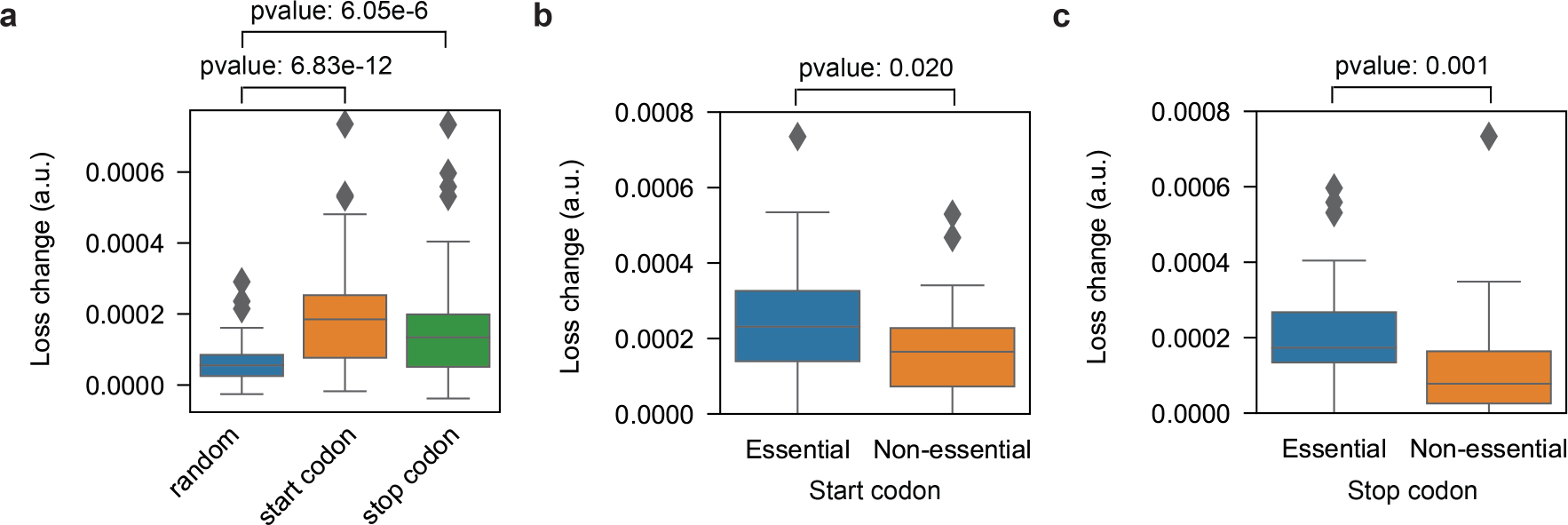
**Model loss changes due to codon mutations**. **a**) Changes in model loss for random 3-nt mutations and mutations in start codon and stop codons of genes in the lambda phage genome (n = 73). **b**) Changes in model loss for mutations in start codons of essential genes (n = 29) and non-essential genes (n = 44). **c**) Changes in model loss for mutations in stop codons of essential genes (n = 29) and non-essential genes (n = 44). For a, b and c, p-values from the Mann-Whitney U test are shown. The central line inside the box represents the median value. The top and bottom borders of the box represent the third (upper) and first (lower) quartiles, respectively.

**Supplementary Figure 3:**
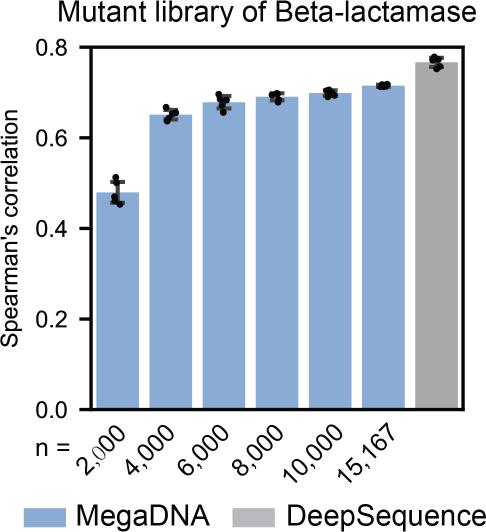
Prediction of mutational effects of beta-lactamase. Spearman correlation of the predicted and reported fitness for mutants from 5-fold cross validation tests are shown. n is the number of training samples for megaDNA. Blue and gray colors represent results from megaDNA and DeepSequence. Error bars denotes standard deviation.

**Supplementary Figure 4:**
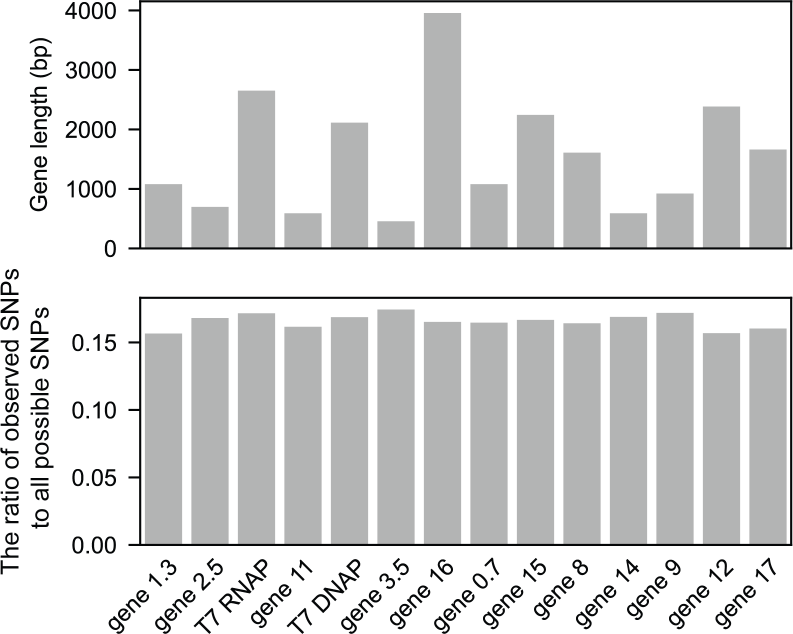
Length and mutation coverage for the genes in T7 bacteriophage genome. Upper: gene lengths. Lower: the ratio of observed SNPs to all possible SNPs per gene.

**Supplementary Figure 5:**
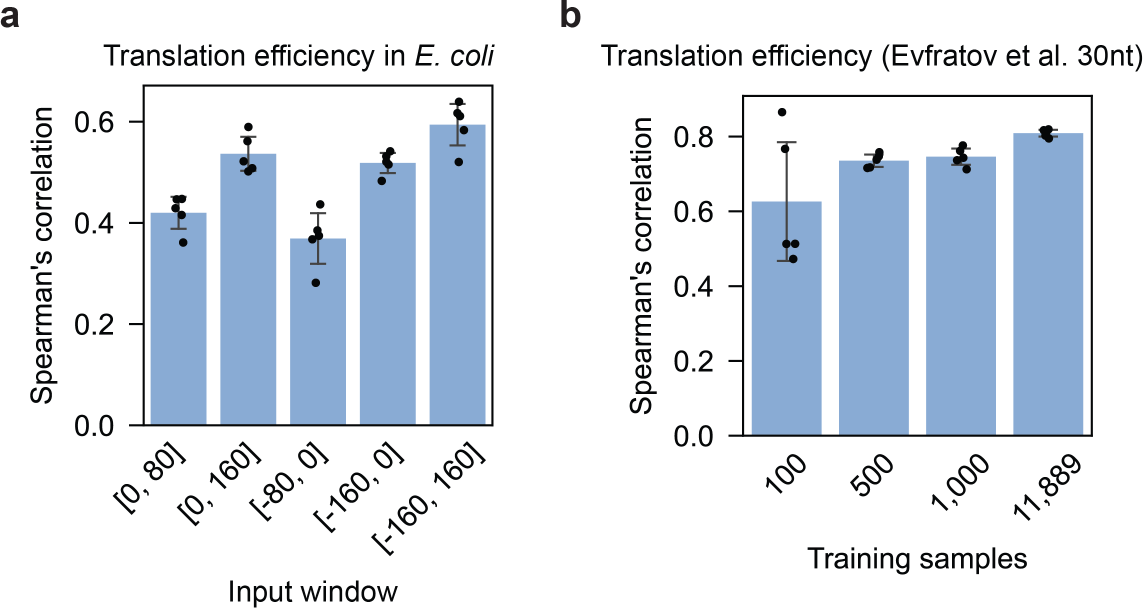
**Impact of window position and training sample size on translation efficiency prediction. a**) Impact of the input window position for the prediction of translation efficiency of endogenous genes in *E. coli*. Positions are reported relative to the start codons. **b**) Effect of training sample number on the prediction performance of 5’UTR activity in the Evfratov et al. dataset. For a and b, results from 5-fold cross validation tests are shown. Error bars denotes standard deviation.

**Supplementary Figure 6:**
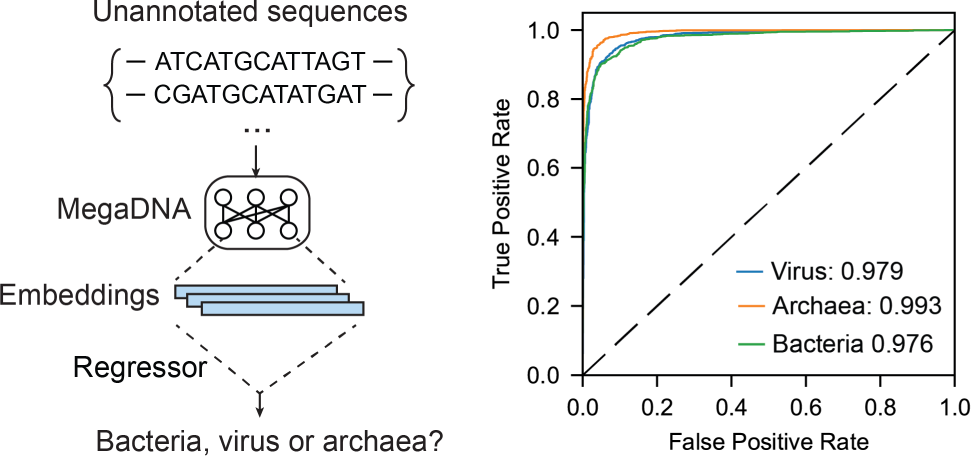
**Taxonomy prediction of unannotated sequences**. Embeddings from the middle layer was used to classify sequences into virus, bacteria, and archaea. The model’s performance was assessed using a 5-fold Stratified K-Fold cross-validation test. The receiver operating characteristic (ROC) curves are shown (n = 5,000 for each category). The mean ROC curve (AUROC) scores from 5-fold cross-validation tests are reported.

**Supplementary Figure 7:**
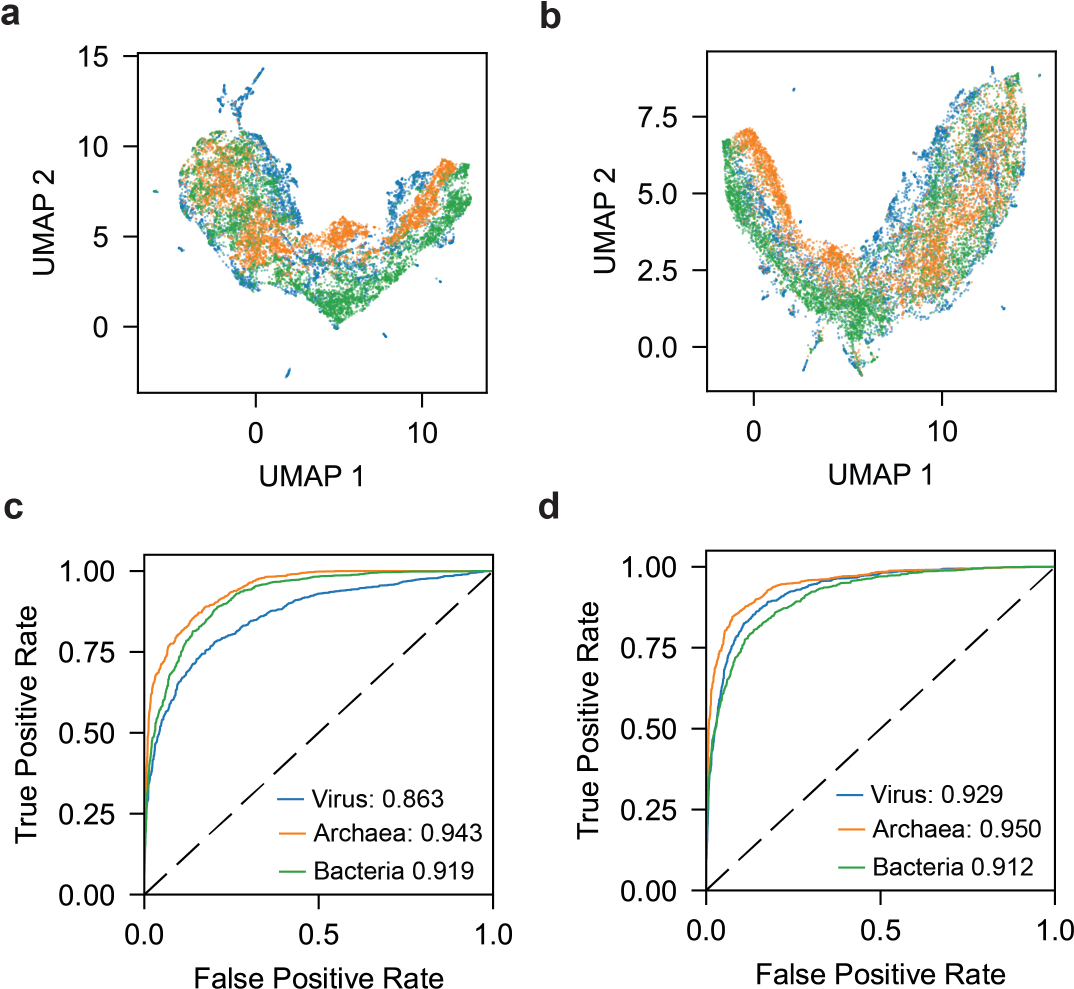
Sequence embedding of different model layers and the taxonomy prediction performance. Sequence embeddings from the local layer (**a**) and global layer (**b**), and their respective taxonomy prediction performances (**c**) and (**d**). The model’s performance was assessed using 5-fold Stratified K-Fold cross-validation tests. The receiver operating characteristic (ROC) curves are shown (n = 5,000 for each category). The mean ROC curve (AUROC) scores from 5-fold cross-validation tests are reported.

**Supplementary Figure 8:**
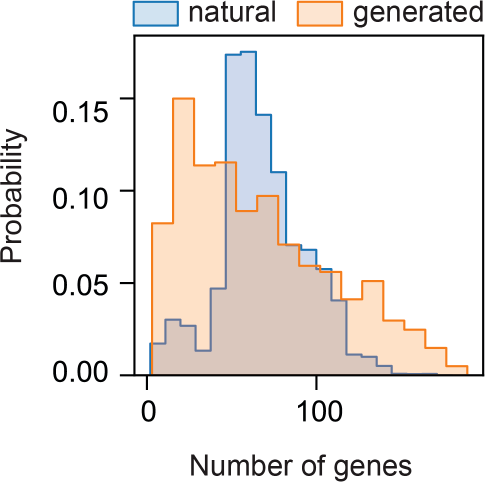
Gene number distributions for the generated sequences and the training dataset. Comparison of the number of predicted genes in generated sequences (n = 607) versus those in the training dataset (n = 99,673)

**Supplementary Figure 9:**
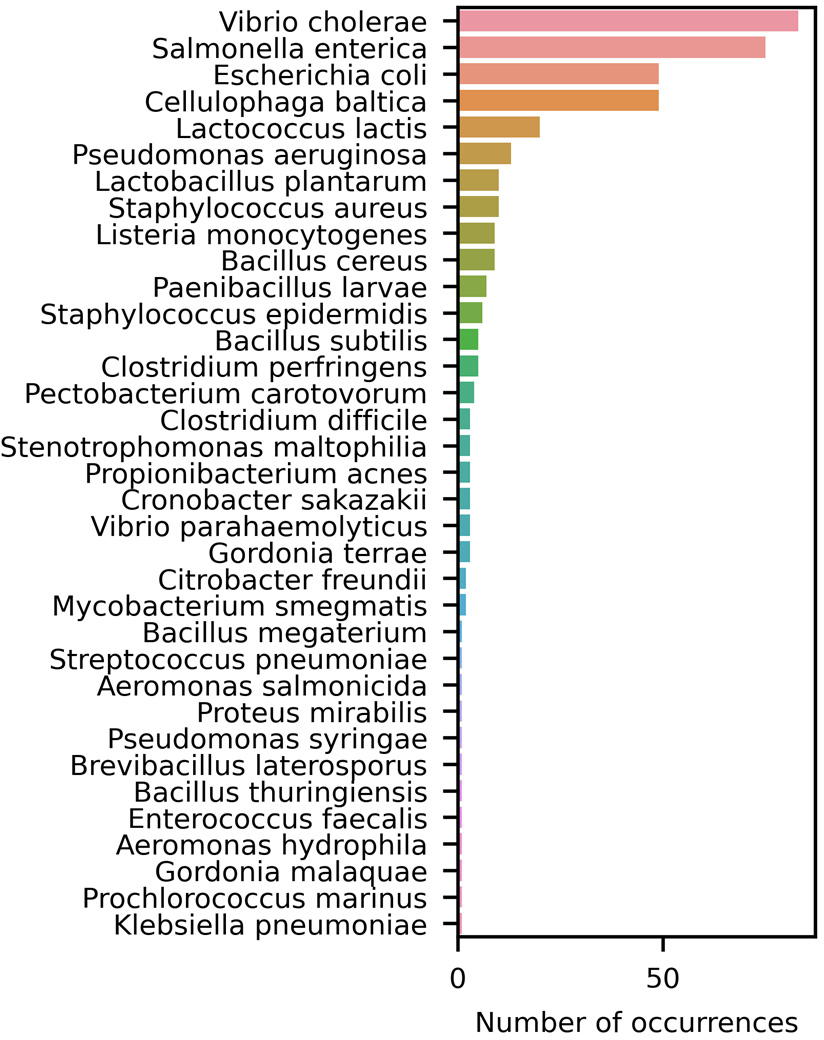
Frequencies of the predicted bacteria hosts for the generated sequences. Host prediction was done using DeepHost^16^ (n = 388). Predictions with a probability greater than 0.95 were used in the analysis.

**Supplementary Figure 10:**
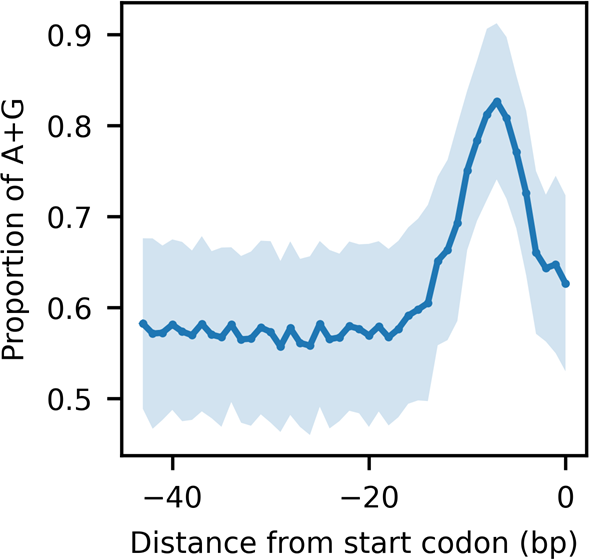
Proportions of adenine (A) and guanine (G) nucleotides preceding the start codon for all the generated sequences. Blue line denotes the mean A+G nucleotides proportion profile for all the generated sequences with a virus score larger than zero (n = 607). The shaded region represents the standard derivation of all profiles.

**Supplementary Figure 11:**
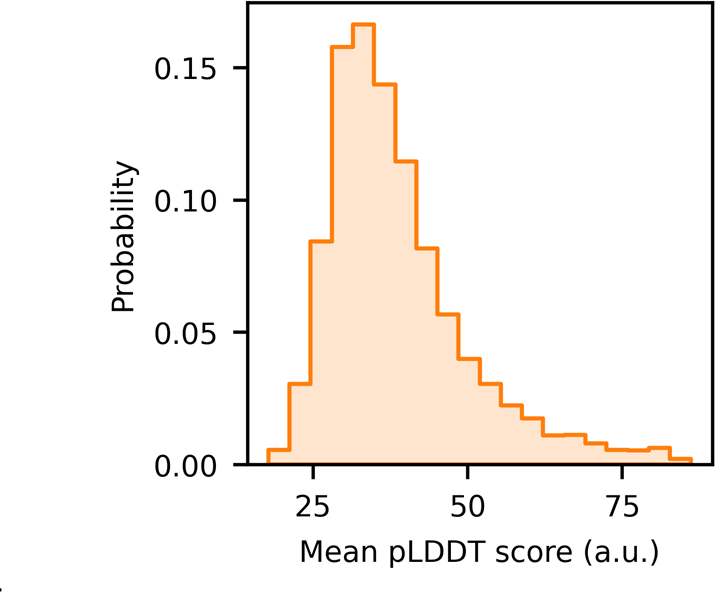
Mean pLDDT scores for proteins derived from the generated sequences. The distribution of mean pLDDT score for a randomly sampled subset of all the generated proteins is shown (sample size: n = 10,000; median value: 36).

**Supplementary Figure 12:**
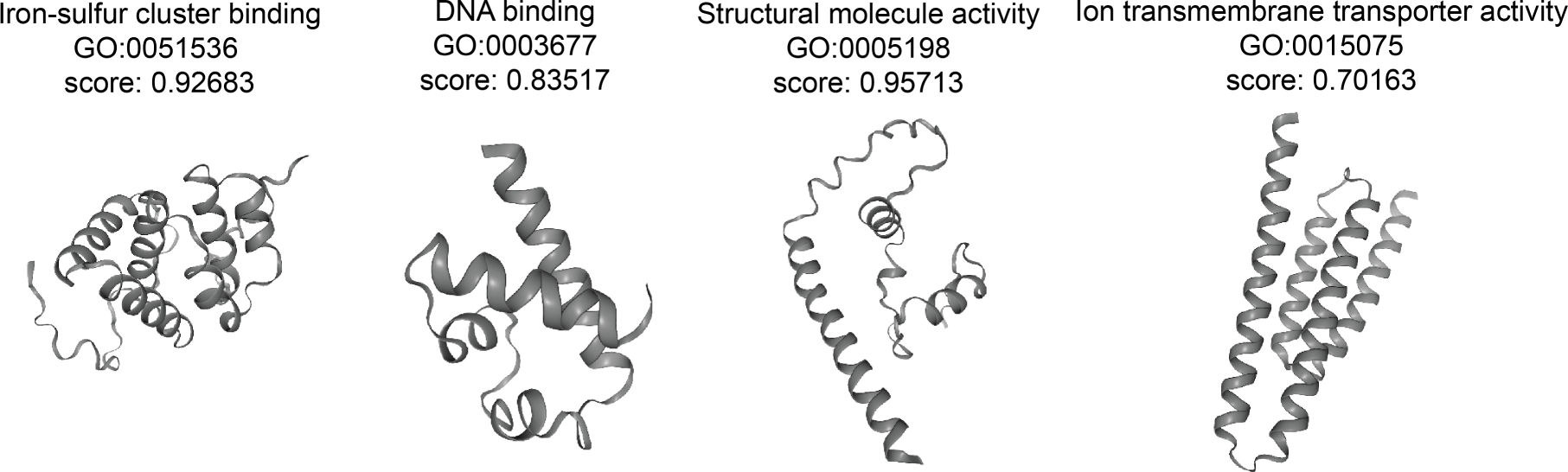
Representative proteins from the generated sequence with predicted functions and structures. The protein structures were predicted using ESMfold^18^ and the functions were annotated using deepFRI^19^. Predicted scores and GO terms from deepFRI are shown.

**Supplementary Table 1:**
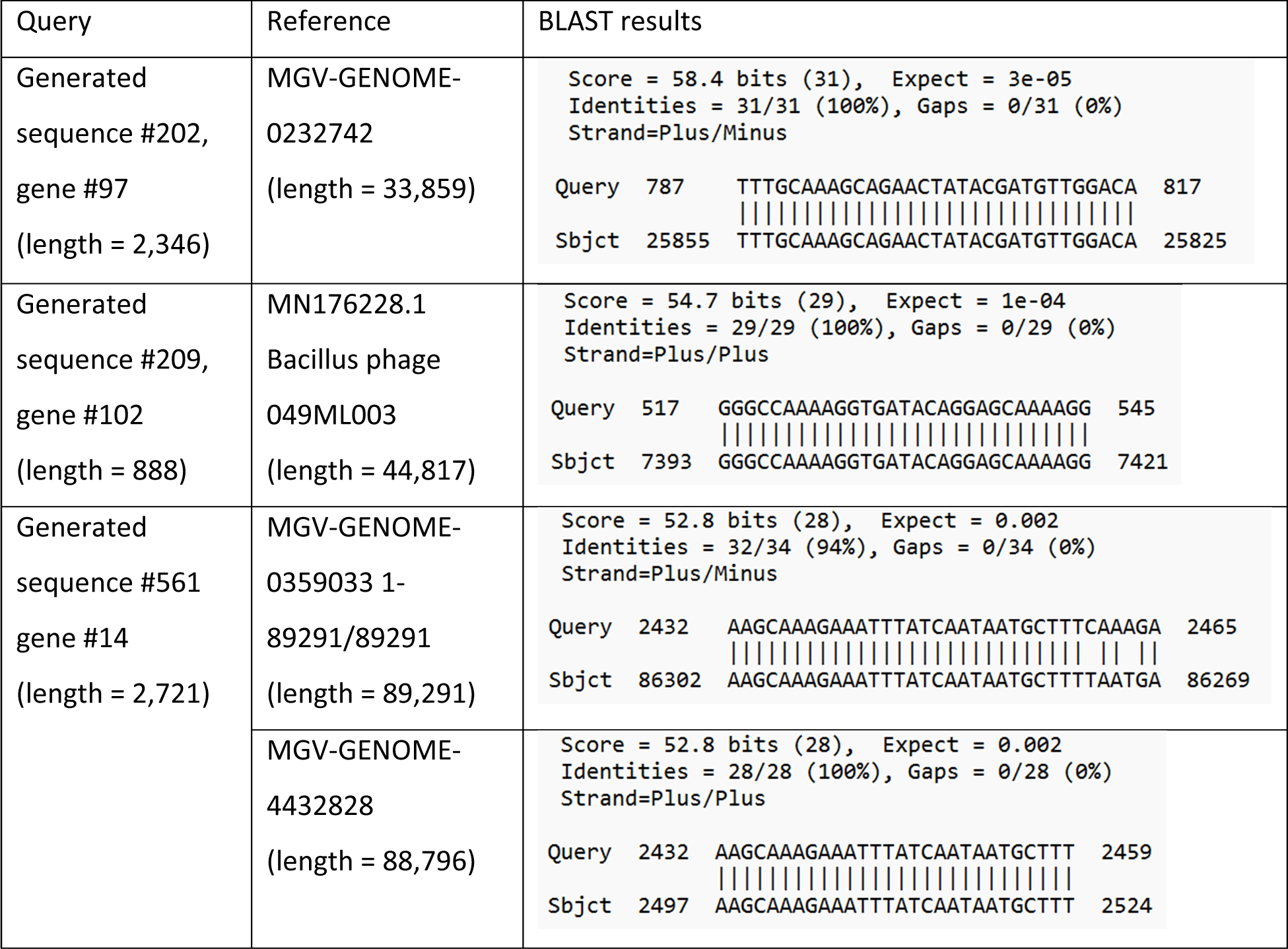
BLAST analysis of the generated genes. We conducted BLAST analysis to compare the generated genes with geNomad markers (n = 343) against the training dataset. Using default settings, we found hits for 3 out of the 343 genes. Only one hit is shown if there are multiple identical matches across reference genomes.

## References

1. Devlin, J., Chang, M.-W., Lee, K. & Toutanova, K. Bert: Pre-training of deep bidirectional transformers for language understanding. arXiv preprint arXiv:1810.04805 (2018).

2. Brown, T. et al. Language models are few-shot learners. Adv Neural Inf Process Syst 33, 1877– 1901 (2020).

3. Ji, Y., Zhou, Z., Liu, H. & Davuluri, R. V. DNABERT: pre-trained Bidirectional Encoder Representations from Transformers model for DNA-language in genome. Bioinformatics 37, 2112–2120 (2021).

4. Dalla-Torre, H. et al. The nucleotide transformer: Building and evaluating robust foundation models for human genomics. bioRxiv 2021–2023 (2023).

5. Benegas, G., Batra, S. S. & Song, Y. S. DNA language models are powerful predictors of genome- wide variant effects. Proceedings of the National Academy of Sciences 120, e2311219120 (2023).

6. Hwang, Y., Cornman, A. L., Kellogg, E. H., Ovchinnikov, S. & Girguis, P. R. Genomic language model predicts protein co-regulation and function. bioRxiv 2023–2024 (2023).

7. Nguyen, E., et al. Hyenadna: Long-range genomic sequence modeling at single nucleotide resolution. *arXiv preprint arXiv:2306.15794* (2023).

8. Yu, L., et al. Megabyte: Predicting million-byte sequences with multiscale transformers. *arXiv preprint arXiv:2305.07185* (2023).

9. Nayfach, S. et al. Metagenomic compendium of 189,680 DNA viruses from the human gut microbiome. Nat Microbiol 6, 960–970 (2021).

10. Camarillo-Guerrero, L. F., Almeida, A., Rangel-Pineros, G., Finn, R. D. & Lawley, T. D. Massive expansion of human gut bacteriophage diversity. Cell 184, 1098–1109 (2021).

11. Piya, D. et al. Systematic and scalable genome-wide essentiality mapping to identify nonessential genes in phages. PLoS Biol 21, e3002416- (2023).

12. Riesselman, A. J., Ingraham, J. B. & Marks, D. S. Deep generative models of genetic variation capture the effects of mutations. Nat Methods 15, 816–822 (2018).

13. Robins, W. P., Faruque, S. M. & Mekalanos, J. J. Coupling mutagenesis and parallel deep sequencing to probe essential residues in a genome or gene. Proceedings of the National Academy of Sciences 110, E848–E857 (2013).

14. Evfratov, S. A. et al. Application of sorting and next generation sequencing to study 5΄-UTR influence on translation efficiency in Escherichia coli. Nucleic Acids Res 45, 3487–3502 (2017).

15. Camargo, A. P. et al. Identification of mobile genetic elements with geNomad. Nat Biotechnol 1– 10 (2023).

16. Ruohan, W., Xianglilan, Z., Jianping, W. & Shuai Cheng, L. I. DeepHost: phage host prediction with convolutional neural network. Brief Bioinform 23, bbab385 (2022).

17. LaFleur, T. L., Hossain, A. & Salis, H. M. Automated model-predictive design of synthetic promoters to control transcriptional profiles in bacteria. Nat Commun 13, 5159 (2022).

18. Lin, Z. et al. Language models of protein sequences at the scale of evolution enable accurate structure prediction. BioRxiv 2022, 500902 (2022).

19. Gligorijević, V. et al. Structure-based protein function prediction using graph convolutional networks. Nat Commun 12, 3168 (2021).

20. Kelsic, E. D. et al. RNA Structural Determinants of Optimal Codons Revealed by MAGE-Seq. Cell Syst 3, 563–571.e6 (2016).

21. Ryu, M.-H. et al. Control of nitrogen fixation in bacteria that associate with cereals. Nat Microbiol 5, 314–330 (2020).

22. Espah Borujeni, A., Zhang, J., Doosthosseini, H., Nielsen, A. A. K. & Voigt, C. A. Genetic circuit characterization by inferring RNA polymerase movement and ribosome usage. Nat Commun 11, 5001 (2020).

23. McInnes, L., Healy, J. & Melville, J. Umap: Uniform manifold approximation and projection for dimension reduction. *arXiv preprint arXiv:1802.03426* (2018).

